# Less cortical complexity in ventromedial prefrontal cortex is associated with a greater preference for risky and immediate rewards

**DOI:** 10.1101/2023.09.12.557368

**Authors:** Fredrik Bergström, Caryn Lerman, Joseph W. Kable

## Abstract

In our everyday lives, we are often faced with situations in which we have to make choices that involve risky or delayed rewards. However, the extent to which we are willing to accept larger risky (over smaller certain) or larger delayed (over smaller immediate) rewards vary across individuals. Here we investigated the relationship between cortical surface complexity in medial prefrontal cortex and individual differences in risky and intertemporal preferences. We found that lower cortical complexity in ventromedial prefrontal cortex (vmPFC) was associated with a greater preference for risky and immediate rewards. In addition to these common structural associations in mPFC, we also found associations between lower cortical complexity and a greater preference for immediate rewards that extended into left dorsomedial prefrontal cortex and right vmPFC. Taken together, the shared association suggests that lower cortical complexity in vmPFC may be a structural marker for individual differences in impulsive behavior.

---

In our daily lives we frequently make decisions that involve risky or delayed rewards. As individuals, we differ in our tendency to accept larger risky (over smaller certain) or larger delayed (over smaller immediate) rewards. An ongoing research question is how such differences in risky or intertemporal preferences are related to differences in brain structure (Kable & Levy, 2015). Cortical surface complexity (Yotter et al., 2011) is a structural property relatively more rooted in genetics and early neurodevelopment (Armstrong, Schleicher, Omran, Curtis, & Zilles, 1995; Kochunov et al., 2010; Zilles, Armstrong, Schleicher, & Kretschmann, 1988), and may therefore be well suited to explain differences in choice behavior. The ventromedial prefrontal cortex (vmPFC) is central to estimating the subjective value of risky or delayed rewards (Bartra, McGuire, & Kable, 2013; Kable & Glimcher, 2009), and lesions to vmPFC disrupt the valuation process (Camille, Griffiths, Vo, Fellows, & Kable, 2011; Fellows & Farah, 2007; Yu, Dana, & Kable, 2022) and can lead to a greater preference for risky and immediate rewards (Mok et al., 2021; Peters & D’Esposito, 2020; Sellitto, Ciaramelli, & Di Pellegrino, 2010). Here we therefore looked for a relationship between cortical complexity in the vmPFC of healthy human participants and preferences for risky and immediate rewards.

Though many brain areas are involved in risky and intertemporal choices (for reviews see Kable & Levy, 2015; Knutson & Huettel, 2015; Mohr et al., 2010; Peters & Büchel, 2011), vmPFC plays a key role in both. Neural activity in vmPFC and ventral striatum (VS) reflects domain-general subjective valuation processes (Bartra et al., 2013; Kable & Glimcher, 2009), and activity in these regions correlates with the value of risky or delayed rewards incorporating individual tolerance levels for risk (Levy, Snell, Nelson, Rustichini, & Glimcher, 2010) or delay (Kable & Glimcher, 2007). Correspondingly, lesions to vmPFC disrupt value maximization and lead to inconsistent choice (Camille et al., 2011; Fellows & Farah, 2007; Yu et al., 2022). Studies on the effects of lesions to vmPFC on risky and intertemporal choice have found mixed results for risky (Clark et al., 2008; Leland & Grafman, 2005; Levens et al., 2014; Manes et al., 2002; Spaniol, Di Muro, & Ciaramelli, 2019; Studer, Manes, Humphreys, Robbins, & Clark, 2015; Weller, Levin, Shiv, & Bechara, 2007) and intertemporal (Fellows & Farah, 2005; Mok et al., 2021; Peters & D’Esposito, 2016; Sellitto et al., 2010) choice separately, but overall tend towards lesions causing greater preference for risky and immediate rewards (for review see, Yu, Kan, & Kable, 2019). Importantly, the two studies that examined the effects of vmPFC lesions on risky and intertemporal choice in the same individuals both found a greater preference for risky and immediate rewards (Mok et al., 2021; Peters & D’Esposito, 2020).

To date, studies linking brain structure to risky and intertemporal preferences have focused on grey matter volume and cortical thickness. The findings are quite varied, though some studies have found relationships with structural properties of medial prefrontal cortex (mPFC). A greater preference for immediate rewards has been positively associated with more grey matter volume (GMV) in mPFC, posterior cingulate cortex (pCC), middle temporal cortex, and entorhinal cortex; less GMV in dorsolateral prefrontal cortex (dlPFC), inferior frontal cortex (IFC), and superior frontal gyrus; both more and less GMV in putamen (Bjork, Momenan, & Hommer, 2009; Cho et al., 2013; Owens et al., 2017; Schwartz et al., 2010); and less cortical surface thickness in mPFC (Bernhardt et al., 2014; Drobetz et al., 2014; Pehlivanova et al., 2018), entorhinal cortex (Lempert et al., 2020), temporal pole and temporoparietal junction (Pehlivanova et al., 2018). A greater preference for risky rewards has been associated with more GMV in amygdalae (Jung, Lee, Lerman, & Kable, 2018), right posterior parietal cortex (PPC) (Gilaie-Dotan et al., 2014; Grubb, Tymula, Gilaie-Dotan, Glimcher, & Levy, 2016), and cerebellum (Quan et al., 2022).

Cortical surface complexity is a measure of how space-filling, self-similar, and convoluted a brain’s surface is (Yotter et al., 2011). High cortical surface complexity has been associated with deeper sulci, higher folding frequency, and more convoluted gyral shape (Im et al., 2006; Jiang et al., 2008; King, Brown, Hwang, Jeon, & George, 2010). The structural properties underlying cortical complexity are deeply rooted in genetics and early neurodevelopment, and may therefore be temporally more stable in adults than grey matter volume (Armstrong et al., 1995; Kochunov et al., 2010; Zilles et al., 1988), but decline with age-related atrophy (Madan & Kensinger, 2016). To date, cortical complexity has mainly been used to study clinical markers (e.g., Hedderich et al., 2020; King et al., 2010; Nenadic et al., 2014, 2017), sex differences (Luders et al., 2006; Luders et al., 2004) and differences in intelligence (Hedderich et al., 2020; Im et al., 2006). However, cortical complexity may also be useful as a structural marker for individual trait differences in decision-making.

Here we investigated the relationship between cortical complexity and individual differences in risky and intertemporal preferences. We focused on mPFC as a region of interest given the role of vmPFC in risky and intertemporal choices and previous work showing that lesions to vmPFC (extending to the greater mPFC area) led to a greater preference for risky and immediate rewards. We find a relationship between lower cortical complexity in left vmPFC and a greater preference for risky and immediate rewards. This finding suggest that cortical folding properties in vmPFC may be a marker for impulsivity.

## Materials and Methods

### Participants

In this study we used a previously published data acquired for the Retraining Neurocognitive Mechanisms of Cancer Risk Behavior (RNMCRB) study. RNMCRB participants were randomized to receive 10 weeks of adaptive cognitive training (Lumosity games) or non-adaptive, untargeted cognitive stimulation (simple computerized video games) and underwent pre- and post-intervention brain scans. The study acquired high-resolution T1-weighted anatomical MRI, resting state fMRI, diffusion tensor imaging, and task-based fMRI during a risky and an intertemporal choice task. Participants were excluded if they had a history of brain injury, a history of psychiatric or substance disorders, current use of psychotropic medication, current use of chewing tobacco, snuff, or smoking cessation products, left-handedness, intellectual disability (< 90 score on Shipley’s Intelligence Quotient (IQ) Test), or had a risk tolerance α < 0.34 or α > 1.32 or discount rate k < 0.0017 or k > 0.077 (to avoid ceiling effects). All study procedures were approved by the Institutional Review Board of the University of Pennsylvania, and all participants provided written informed consent. The main trial outcomes have been described in a previous report, but found no effect of cognitive training relative to active control on brain activity, decision-making, or cognitive performance (Kable et al., 2017).

For this study we used 166 anatomical high-resolution T1-weighted images from the pre-intervention (baseline) session from Kable et al. (2017). We excluded nine participants for which it was impossible to reliably estimate preference for risky or delayed rewards because of one-sided choices. Additionally, we quality-checked the cortical complexity image quality in CAT12 by performing overall correlations between all participants (to check sample homogeneity) and found that five participants had an overall correlation value less than two standard deviations from the median. At closer inspection, one of the participants had an artifact in the left vmPFC and was therefore excluded. We therefore used 156 participants (90/66 males/females; age, M = 25, SD = 6 years; IQ, M = 111, SD = 7; delay preference (log10k), M = −1.74, SD = 0.36; risk preference (log10α), M = −0.20, SD = 0.17) in our analyses.

### Risky choice task

In the risky choice task, participants had to make a choice between receiving a smaller but certain reward (i.e., 100% probability to receive $20) or a larger but riskier reward (e.g., 47% probability to receive $84) on 120 trials. The certain reward was always fixed at $20 on all trials, while the probability and risky reward amount varied across trials. Each trial began by presenting probability and reward amount for the risky alternative (the fixed certain option was never displayed). Participants had 4 s to accept or reject the risky alternative, after which a marker indicating their choice (a checkmark if the risky alternative was accepted, and “X” if rejected) appeared for 1 s. Based on each participant’s choices, we estimated their individual risk tolerance (α) by fitting a logistic regression to their choice data. The subjective value (SV) of the choice options was assumed to follow a power utility function:

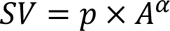

where p is the probability of winning amount A, and α is a risk tolerance parameter that varies across participants. For the risky option, there is always a 1 - p chance of not winning anything. Higher α indicates higher risk tolerance or lower risk aversion.

### Intertemporal choice task

In the intertemporal choice task, participants had to make a choice between receiving a smaller but immediate reward (i.e., receiving $20 today) or a larger but delayed reward (e.g., receiving $40 in 31 days). The immediate reward was always fixed at $20 on all trials, while the delay time and delayed reward amount varied across trials. Each trial began by presenting the delay time and reward amount of the delayed option (the fixed immediate option was never displayed). The participants had 4 s to accept or reject the delayed alternative, after which a marker indicating the choice (a checkmark if the delayed alternative was accepted, and “X” if rejected) appeared for 1 s. Based on each participant’s choices, we estimated their individual delay discount rate (k) by fitting a logistic regression to their choice data. The subjective value (SV) of the choice options was assumed to follow hyperbolic discounting:

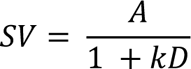

where A is the amount of the delayed option, D is the delay time until receiving the reward (for immediate choice, D = 0), and k is a discount rate parameter that varies across participants. Higher values of k indicate less tolerance for delay or more impatience.

### MRI acquisition

High-resolution T1-weighted anatomical images were acquired by a Siemens 3T Trio scanner (Siemens, Erlangen, Germany) using a magnetization-prepared rapid gradient echo (MPRAGE) sequence [repetition time (TR) = 1630 ms, echo time (TE) = 3.11 ms, voxel size = 0.94 x 0.94 x 1.0 mm, 160 axial slices, 192 x 256 matrix].

### Preprocessing of MRI data

We used the default processing pipeline of the Computational Anatomy Toolbox (CAT12; Gaser & Dahnke, 2016) for SPM12 (Welcome Trust Centre for Neuroimaging, London, UK), on Matlab R2019a (Mathworks, Inc., Sherborn, MA, USA), to process the anatomical T1-weighted images, and extract local cortical surface complexity (i.e., fractal dimension) values.

For voxel-based preprocessing, T1-weighted images underwent spatial adaptive non-local means (SANLM) denoising filter (Martı & Manjo, 2010), were internally resampled, bias corrected, and affine-registered, followed by standard SPM unified tissue segmentation into grey matter, white matter, and cerebral spinal fluid (Ashburner & Friston, 2005). The resulting brain images were then skull-stripped, parcellated into left and right hemispheres, subcortical areas, and cerebellum; subject to local intensity transformation of all tissue classes to reduce effects of higher grey matter intensities before the final Adaptive Maximum A Posteriori (AMAP) tissue segmentation (Rajapakse, Giedd, & Rapoport, 1997), and refined by a partial volume estimation (Tohka, Zijdenbos, & Evans, 2004).

For surface-based preprocessing, a projection-based thickness method was used to estimate and reconstruct cortical thickness of the central surface (Dahnke, Yotter, & Gaser, 2013), after which topological correction was applied to repair defects with spherical harmonics (Yotter et al., 2011), and surface refinement. The final central surface mesh was used to estimate local cortical surface complexity (i.e., fractal dimension) values based on spherical harmonic reconstructions (Yotter et al., 2011). The cortical complexity maps were spatially registered to a surface template, resampled to 1.5 mm^3^, and spatially smoothed with a 20 mm Gaussian FWHM kernel (as recommended in the CAT12 manual; http://www.neuro.uni-jena.de/cat12/CAT12-Manual.pdf) before statistical analysis.

### Statistical analysis of fMRI data

First, we performed a restricted vertex-wise region of interest analysis within the mPFC. Second, we performed an exploratory whole cortical surface analysis. For our region of interest analysis, we created a mask by combining anatomical mPFC areas (Fo1, Fo2, Fo3, Fp1, Fp2) from The Anatomy Toolbox (Eickhoff et al., 2007, 2005), and a 30 mm sphere centered on MNI coordinates XYZ = [−1 46 −7] (the domain general valuation system; Bartra et al., 2013) to “fill-in” some small gaps between the anatomical vmPFC atlas regions. This mPFC mask was transformed from volumetric to surface template MNI space with CAT12. For all analyses, we used mass-univariate multiple linear regressions with log10-transformed α (risky preferences) and k (intertemporal preferences) as variables of interest, while controlling for demographic variables (i.e., age, sex, and IQ). Non-parametric statistical t-maps were adjusted with threshold-free cluster enhancement (TFCE) and FDR (q < 0.05) corrected based on 10,000 permutations using the TFCE toolbox (http://www.neuro.uni-jena.de/tfce/). We used a conjunction analysis to look for regions linked to both a greater preference for risky rewards (higher log10α) and a greater preference for immediate rewards (higher log10k). For a vertex to qualify as significant in our conjunction analysis, it had to have a significant association with both risky (log10α) and intertemporal (log10k) preferences, independently (Nichols, Brett, Andersson, Wager, & Poline, 2005).

## Results

We found that cortical complexity in our medial frontal region of interest was associated with preferences for risky and immediate rewards. The conjunction analysis identified a region in left vmPFC, extending into orbitofrontal cortex (OFC), where less cortical surface complexity was associated with both a greater preference for risky and a greater preference for immediate rewards (Figure 1A). This association was stronger more anteriorly and bordering frontopolar cortex. When analyzing risky preferences, there were no distinct associations in mPFC (Figure1B) beyond the region of left vmPFC identified in the conjunction analysis, where there was a negative association between cortical surface complexity and preference for risky rewards (peak MNI = [-7 46 −10], TFCE value = 21784, k = 2607). When analyzing intertemporal preferences, however, the negative association between cortical complexity and preferences for immediate rewards extended beyond the left vmPFC region identified in the conjunction (Figure 1C), and included dorsomedial PFC (peak MNI = [-7 25 23], TFCE value = 20536, k = 6042) and right vmPFC (peak MNI = [15 15 −12], TFCE value = 3640, k = 369).

**Figure 1.**
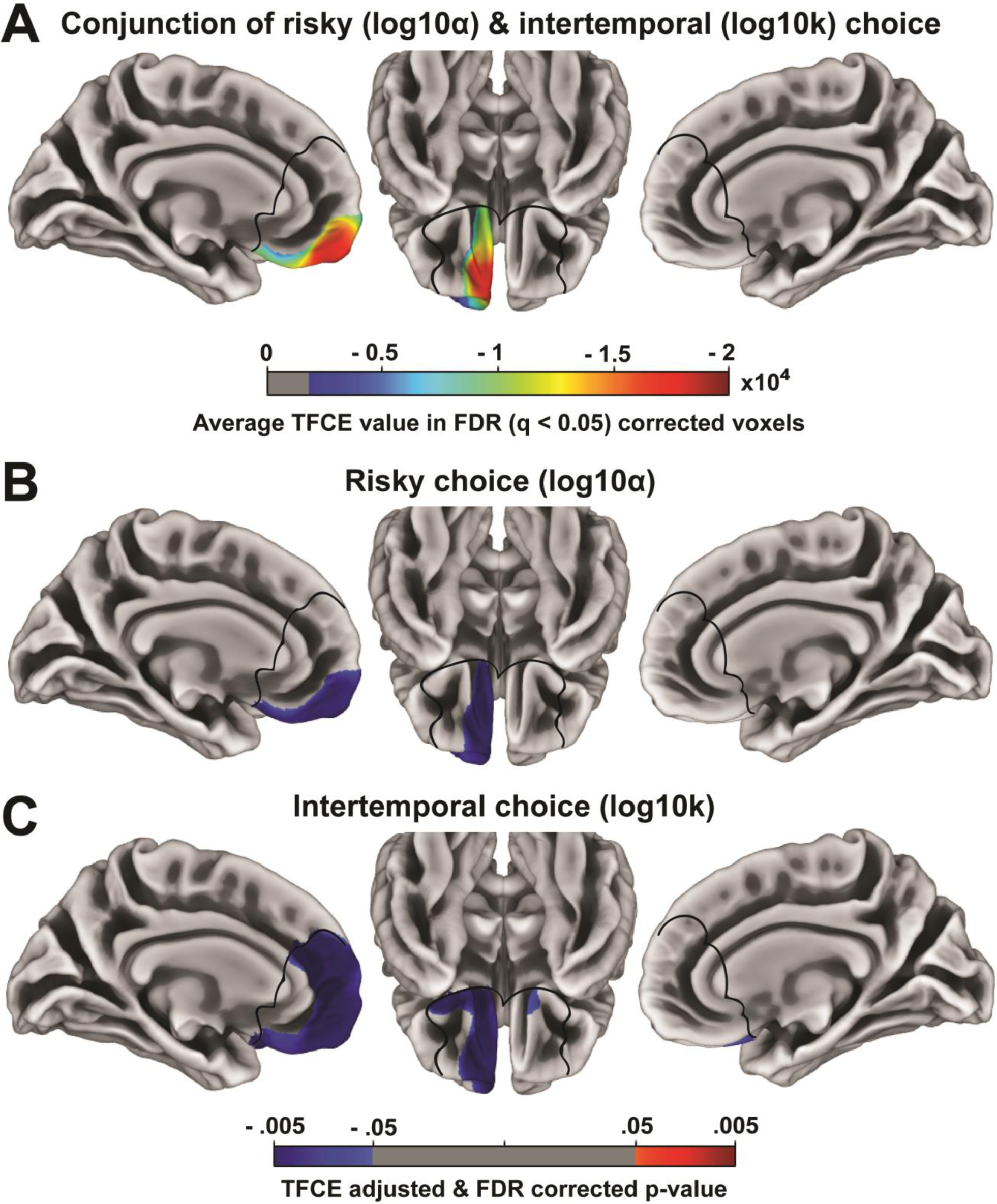
Region of interest results within mPFC. (**A**) Shows the average TFCE value for vertices in which less cortical complexity was associated with a greater preference for risky and immediate rewards. (**B**) Shows associations between cortical complexity and risky choice preferences. (**C**) Shows associations between cortical complexity and intertemporal choice preferences. The black lines outline the mPFC mask used to restrict the analyses to our region of interest. All results are threshold-free cluster-enhancement (TFCE) adjusted and FDR (q < 0.05) corrected.

To identify any regions where cortical complexity was associated with risky or intertemporal preferences beyond our region of interest, we also performed an exploratory whole cortical surface analysis. No results survived FDR (q < 0.05) correction. To potentially generate hypotheses for future studies, we present uncorrected results in Extended Figure 1-1.

**Extended Figure 1-1.**
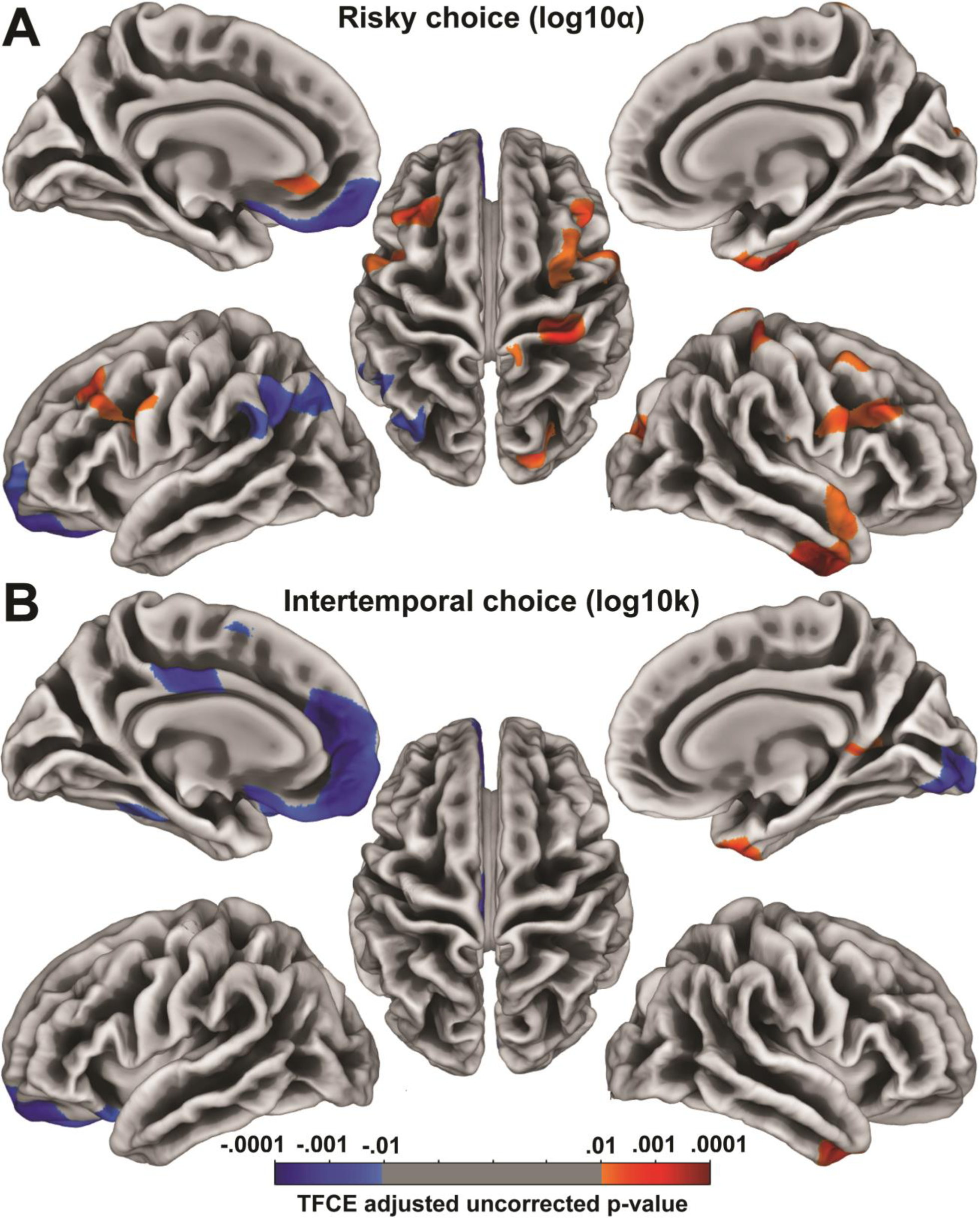
Uncorrected whole cortical surface results. (**A**) Shows associations between cortical complexity and risky choice preferences (log10α). (**B**) Shows associations between cortical complexity and intertemporal choice preferences (log10k). All results are threshold-free cluster-enhancement (TFCE) adjusted and uncorrected.

## Discussion

Here we investigated the relationship between local cortical surface complexity in mPFC and individual differences in preferences for risky and immediate rewards. We found that less cortical complexity in left vmPFC, extending in to orbitofrontal cortex and frontopolar cortex, was associated with a greater preference for both risky and immediate rewards. The associations between cortical complexity and intertemporal choice also extended beyond this region, such that a preference for immediate rewards was associated with less cortical complexity in left dmPFC and right vmPFC. Taken together, these findings suggest that cortical complexity in vmPFC is a structural marker for individual trait differences in risky and intertemporal choice behavior – and possibly a marker for general impulsivity.

That less cortical complexity in vmPFC is associated with a greater preference for both risky and immediate rewards is a novel finding consistent with our expectations based on the neuropsychological literature. Specifically, individuals with vmPFC lesions have a greater preference for risky and immediate rewards (Mok et al., 2021; Peters & D’Esposito, 2020). However, large lesions are not as informative as to whether shared or distinct areas in vmPFC are responsible for these effects on risky and intertemporal choices. Here we were able to show that cortical folding properties in the same area of left vmPFC are associated with both risky and intertemporal preferences in healthy participants.

Our results also contribute to a growing understanding of functional subdivisions within vmPFC. Interestingly, the region where cortical complexity is associated with risky and intertemporal preferences is anterior and ventral to area of vmPFC where activity reflects the subjective value of choice options across domains (Bartra et al., 2013), including in risky, intertemporal, and effortful choice (Seaman et al., 2018). Several studies found that lesions to more ventral areas of vmPFC and OFC caused a stronger preference for immediate rewards (Mok et al., 2021; Peters & D’Esposito, 2016; Sellitto et al., 2010), while the Fellows & Farah (2005) study that found no effect on intertemporal preferences included individuals with more dorsal vmPFC lesions. Moreover, in a large study of adolescents and young adults, a greater preference for immediate rewards was associated with reduced cortical thickness in similar region of vmPFC/OFC to the one identified here (Pehlivanova et al., 2018). It is less clear whether there is a similar ventral-dorsal difference for risky preferences because the lesion loci and tasks vary widely across studies (Clark et al., 2008; Leland & Grafman, 2005; Levens et al., 2014; Manes et al., 2002; Spaniol et al., 2019; Studer et al., 2015; Weller et al., 2007). Nevertheless, the two studies using larger groups with focal lesions that overlap with the more anterior and ventral region of vmPFC/OFC found a greater preference for risky rewards (Clark et al., 2008; Studer et al., 2015). One possibility is therefore that the ventral and anterior aspects of vmPFC/OFC have a different function than mid-vmPFC, which seems likely given the hierarchical functional organization of the PFC and more granular structural organization from mPFC to OFC/frontopolar cortex (for review see, Wallis, 2012).

Reduced cortical complexity in vmPFC may be a structural marker for individual trait differences in impulsivity, given that it is associated with a greater preference for both larger risky over smaller certain rewards and smaller immediate over larger delayed rewards. However, the idea of a unified concept of impulsivity is problematic because just as there are groups of people who are risk-seeking and impatient, there are also groups who are risk-averse and impatient, and groups who have an equal tolerance for risk and delay (for critical review see Green & Myerson, 2013). Furthermore, people tend to have different preferences for different rewards such as money or health outcomes (Green & Myerson, 2013). As such, it is likely that the neural networks underpinning our risky and intertemporal preferences consist of both shared and distinct associations with brain structure. Although we have pinpointed a shared association in vmPFC with risky and intertemporal preferences, there are likely other distinct associations that contribute to divergences between risky and intertemporal preferences. However, our findings do imply that, all else equal, if there is a lesion or other degradation of cortical complexity in vmPFC (e.g., from clinical disorders, drug use, or age), then it would lead to more risk-seeking and impatient behavior.

We also found that the association between cortical complexity and intertemporal choice preferences extended beyond left vmPFC, most notably into the left dmPFC. Although we can only speculate about the functional significance of the association in dmPFC, neural activity in the same dmPFC has previously been associated with the subjective value during intertemporal choice (Kable & Glimcher, 2007). Similarly, structural studies have found that a greater preference for immediate rewards is associated with more GMV (Cho et al., 2013) and more cortical thickness (Bernhardt et al., 2014) in dmPFC.

In conclusion, we found that less cortical surface complexity in the vmPFC was associated with a greater preference for both risky and immediate rewards. This association with risky and intertemporal preferences is related to cortical folding properties that depend relatively more on genetics and early neurodevelopment and may reflect individual differences in impulsivity. Our result is consistent with previous neuropsychological findings that vmPFC lesions lead to a greater preference for risky and immediate rewards. Future work should further consider the relationship between cortical complexity (and other structural properties beyond GMV and cortical thickness) and choice behavior.

## Acknowledgements

This work was supported by Fundação para a Ciência e Tecnologia (CEECIND/03661/2017 to F.B.); and National Cancer Institute Grants (R01-CA-170297 to J.W.K. and C.L., and R35-CA-197461 to C.L.).

## Notes

https://doi.org/10.18112/openneuro.ds002843.v1.0.1

### Competing Interest Statement

The authors have declared no competing interest.

